# Oncogenic Ras, Yki and Notch signals converge to confer clone competitiveness through Upd2

**DOI:** 10.1101/2023.04.05.535774

**Authors:** Ying Wang, Jingjing He, Mingxi Deng, Yan Yan

## Abstract

It has long been proposed that cell competition functions to remove precancerous clones. A classical model is the removal of polarity-deficient clones such as the *scribble* (*scrib*) mutant clones in *Drosophila* imaginal discs. The activation of Ras, Yki or Notch signaling robustly reverses the *scrib* mutant clonal fate from elimination to tumorous growth. Using single-cell transcriptomics techniques to profile wing imaginal discs harboring the *scrib* mutant clones in combination with different signals, we found that a critical converging point downstream of Ras, Yki and Notch signals is the upregulation of Upd2, which is necessary to promote tumorous growth. Unexpectedly, while Upd2 is not required for cell survival *per se*, Upd2-deficient clones are efficiently wiped out from epithelia, indicating that Upd2 is a previously unrecognized cell competition factor.

## Introduction

Cell competition refers to the phenomenon that viable cell clones are selectively eliminated when they encounter neighbors of higher fitness (*1*–*4*). Originally discovered during the study of Minute mosaic clones in *Drosophila* wing discs (*5*), cell competition is now recognized as perhaps a common surveillance mechanism to eliminate various types of less-fit cells across metazoan species (*3*). Among its many functions in maintaining organism fitness, a tumor-suppressive role of cell competition, which leads to the elimination of precancerous cell clones, was found in both *Drosophila* and humans (*3*, *6*).

In *Drosophila*, the *scribble* (*scrib*) gene encodes a conserved apicobasal cell polarity gene (*7*, *8*). When the larval imaginal discs and optic lobes are homozygous mutant for the *scrib* gene, these tissues grow into amorphous and lethal tumors (*8*). However, when the *scrib* mutant cells are generated as mosaic clones surrounded by wild type cells in imaginal discs, the *scrib* mutant clones are efficiently eliminated and the larvae are free of tumor burden (*9*, *10*). The elimination of the *scrib* mutant clones is generally regarded as a process of cell competition. For example, Sas and PTP10D were identified as a ligand-receptor pair that mediates the recognition between wild type cells and the *scrib* mutant clones, which then triggers cell competition (*11*). However, a recent study demonstrates that the elimination of the *scrib* mutant clones is caused by the exposure of apical TNFR to circulating TNF/Eiger associated with loss of epithelial integrity (*12*). This study suggests that the elimination of the *scrib* mutant clones is independent of influences from neighboring cells (*12*). Notably, it has long been recognized that the activation of a few oncogenic signals, including the co-expression of Ras^V12^, Yki^S168A^, or Notch intracellular domain (N^Act^) constructs, can rescue the *scrib* mutant clones from elimination and promote tumor formation (*9*, *10*, *13*). Whether Ras^V12^, Yki^S168A^ and N^Act^ signals converge at critical downstream points to reverse the *scrib* mutant clone fate from elimination to tumorous growth remain unclear.

## Results

### Single-cell RNA sequencing of wing discs harboring the *scrib* mutant clones in combination with Ras^V12^, Yki^S168A^ or N^Act^ signals

We first generated the *scrib* mutant clones with activated Ras, Yki or Notch signals using MARCM system in wing imaginal discs (Figure 1A-E). The control MARCM GFP+ clones occupy around 30% of the whole disc area at 36 hours after clone induction (ACI) and around 40% of the disc area at 60 hours ACI (Figure 1A, 1A’, 1F, 1F’). In contrast, the *scrib* mutant clones occupy around 20% of the whole disc area at 36 hours ACI and 10% of the whole disc area at the 60 hours ACI (Figure 1B, 1B’, 1F, 1F’). When Ras^V12^, Yki^S168A^ or N^Act^ is coexpressed in the *scrib* mutant clones, the GFP+ clones occupy around 30% of the whole disc area at 36 hours ACI (Figure 1C-E). By 60 hours ACI, the *scrib* mutant clones co-expressing Ras^V12^, Yki^S168A^ or N^Act^ start to exhibit tumor-like morphology and occupy around 30%-40% of the whole disc area (Figure 1C’-F’). Therefore, our experimental setting is able to fully recapitulate the reported phenomena of the *scrib* mutant clone elimination and the *scrib* mutant clonal rescue by Ras^V12^, Yki^S168A^ or N^Act^ signals.

**Figure 1.**
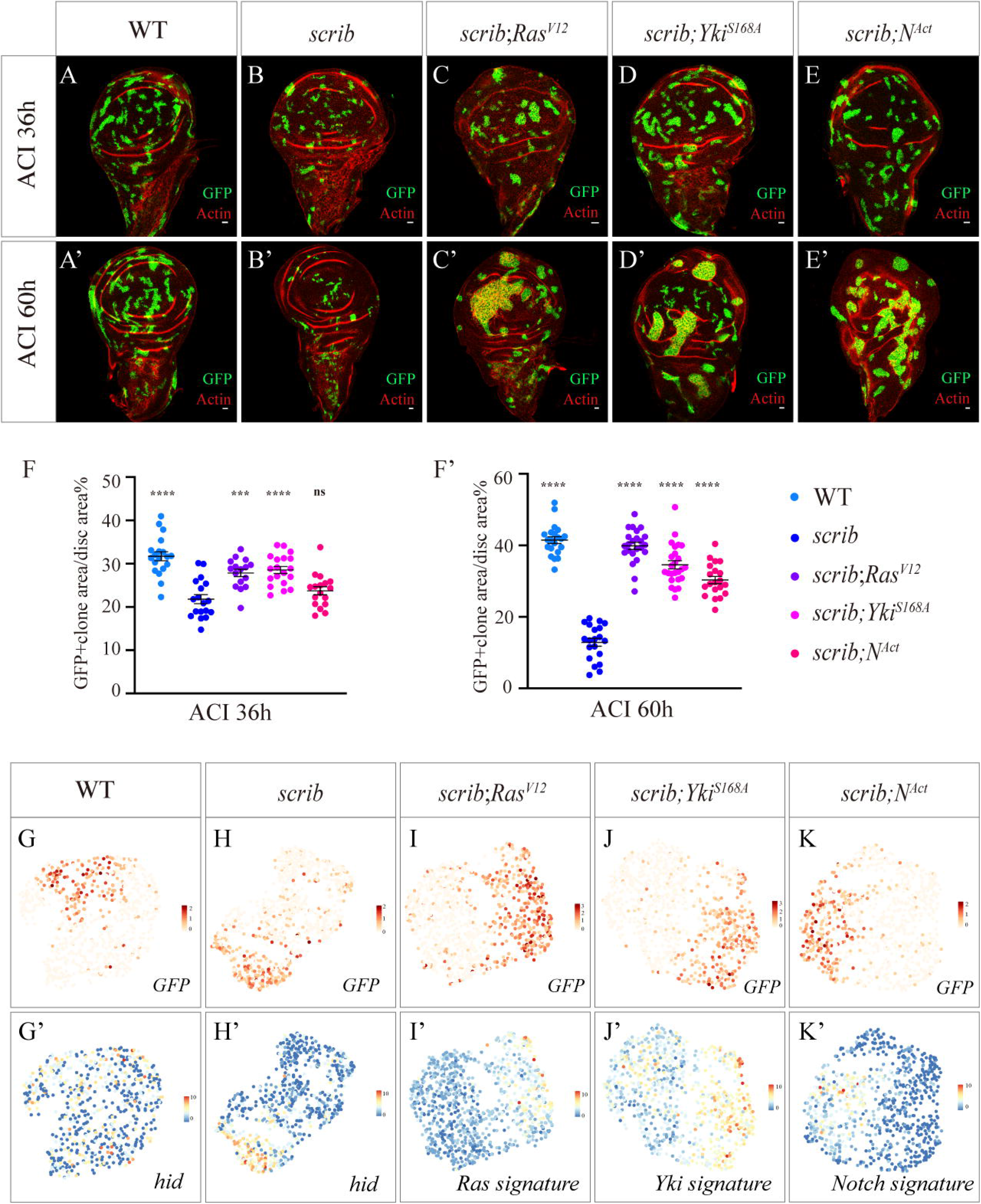
Single-cell RNA sequencing analysis of wing discs harboring the *scrib* mutant clones in combination with Ras^V12^, Yki^S168A^ or N^Act^ signals. (A-E) Confocal images of wing imaginal discs harboring MARCM clones of indicated genotypes inspected at 36 hours after clone induction (ACI) (A-E) and 60 hours ACI (A’-E’). The MARCM clones are marked by the presence of GFP (green) and imaginal discs are stained with phalloidin to visualize cell outline (red). Scale bar: 10μm. (F) Quantification of the percentage of MARCM GFP+ clone area in wing discs. ns p>0.05, *** p<0.001, **** p<0.0001. One-way ANOVA test. Error bars represent mean with SEM. (G-K) UMAP projection of single cells from wing discs harboring MARCM clones of indicated genotypes. The cells are clustered using genes correlated with GFP gene expression for visualization. (G-K) are colored by the expression level of GFP. (G’-K’) are colored as the expression score of *hid* (G’-H’) or respective gene sets (I’-K’).

Using this experimental setting, we generated single-cell RNA transcriptomics datasets of wing discs carrying GFP+ clones of these genotypes: control, *scrib*^-/-^, *scrib*^-/-^ *Ras^V12^*, *scrib*^-/-^ *Yki^S168A^* and *scrib*^-/-^ *N^Act^* at 36 hours ACI. We chose to profile wing discs at 36 hours ACI because the clone size and morphology are comparable across all five experimental groups at this stage (Figure 1A-E). In another words, cell competition, which is a hurdle that these oncogenic signals need to overcome, must still be an on-going process at this early stage. For each experimental group we captured from around 4,500 to 10,000 high-quality cells and detected around 3000 genes per cell on average (Supplemental Table 1). In each group, around 3% ~ 6% of captured cells express GFP (Supplemental Table 1), which is lower than the actual percentage of GFP+ cells (20%~30%) in these wing discs. This is likely due to the dropout phenomenon commonly observed in all single-cell RNA sequencing experiments. These cells largely fall into two clusters, wing disc epithelial cells (ECs) and myoblasts, in the UMAP projection (Supplemental Figure 1A). For the rest of this study, we only focus on the wing disc epithelial cell cluster. Notably, across all experimental groups, the proximal-distal (PD) patterning genes such as *nub* and *tsh* remain the primary source for cell clustering, consistent with previous reports (Supplemental Figure 1B) (*14*, *15*). In order to better visualize the GFP+ clonal signature, we randomly selected 500 GFP-cells from each group to balance GFP+ and GFP-cell number. The UMAP projection shows that PD patterning genes are still the main source for clustering (Supplemental Figure 2). We then projected these cells using genes correlated with GFP expression (Supplemental Table 2) to better visualize the differences between GFP+ and GFP-cells (Figure 1G-K). For example, in comparison with GFP-cells, the *scrib* mutant clones express higher level of apoptotic genes such as *hid*, consistent with their loser cell status (Figure 1G’-H’). Previous studies have shown that the expression of Ras^V12^ induces profound transcriptomic changes in imaginal discs (*16*, *17*). We extracted a Ras signature gene set composed of 138 genes significantly upregulated in the *scrib*^-/-^ *Ras^V12^* mutant tumors in comparison with control from published datasets (*17*) (Supplemental Table 3). We mapped the expression of Ras signature gene set to our data and found that the *scrib*^-/-^ *Ras^V12^* clones indeed exhibit higher expression level of this gene set in comparison with GFP-cells (Figure 1I’). Similarly, we mapped an Yki signature gene set composed of 52 genes to our data (*18*) (Supplemental Table 3) and found that the *scrib*^-/-^ *Yki^S168A^* clones exhibit higher expression level of Yki target genes in comparison with GFP-cells (Figure 1J’). The *scrib*^-/-^ *N^Act^* clones express higher level of *the enhancer of split* gene complex, which are well-known Notch signaling targets (*19*) (Figure 1K’). Together, these data suggest that our single-cell RNA sequencing experiments are able to capture the expected signatures of each clonal type.

### The *scrib*^-/-^ *Ras^V12^*, *scrib*^-/-^ *Yki^S168A^* and *scrib*^-/-^ *N^Act^* clones exhibit transcriptomic signatures indicative of largely similar cell proliferation, cell death and cell growth states

Compared with the *scrib* mutant clones that undergo elimination, the *scrib*^-/-^ *Ras^V12^*, *scrib*^-/-^ *Yki^S168A^* and *scrib*^-/-^ *N^Act^* clones are phenotypically similar in that these clones can evade cell death and develop into tumors. With the single-cell RNA sequencing data, we can now examine the similarity and differences of these clones at the molecular level. We started by examining gene sets which are indicative of cell proliferation, death and growth states, which are the major parameters determining the *scrib* mutant clonal growth outcome.

We have previously identified a cell proliferation gene set composed of 47 genes through gene expression correlation analysis of wing disc single cells (*14*) (Supplemental Table 3). This gene set contains genes such as *CycE, cdc25/string, PCNA* and *Mcm* family genes, which are likely co-regulated by the E2F1 factor (*20*). We found that the GFP+ cells exhibit lower expression level of the cell proliferation gene set except in the control and N^Act^ groups (Figure 2A’-2E’). We quantified a proliferation index of these clones using phospho-Histone H3 (pH3) staining as a marker (Figure 2F and 2F’). We observed that the proliferation index of the *scrib* mutant clones is significantly lower than that of control clones, and more importantly, the proliferation index of the *scrib* mutant clones does not increase when Ras^V12^, Yki^S168A^ or N^Act^ is co-expressed in the clones, indicating that Ras^V12^, Yki^S168A^ and N^Act^ signals likely do not enhance cell proliferation capacity at this stage.

**Figure 2.**
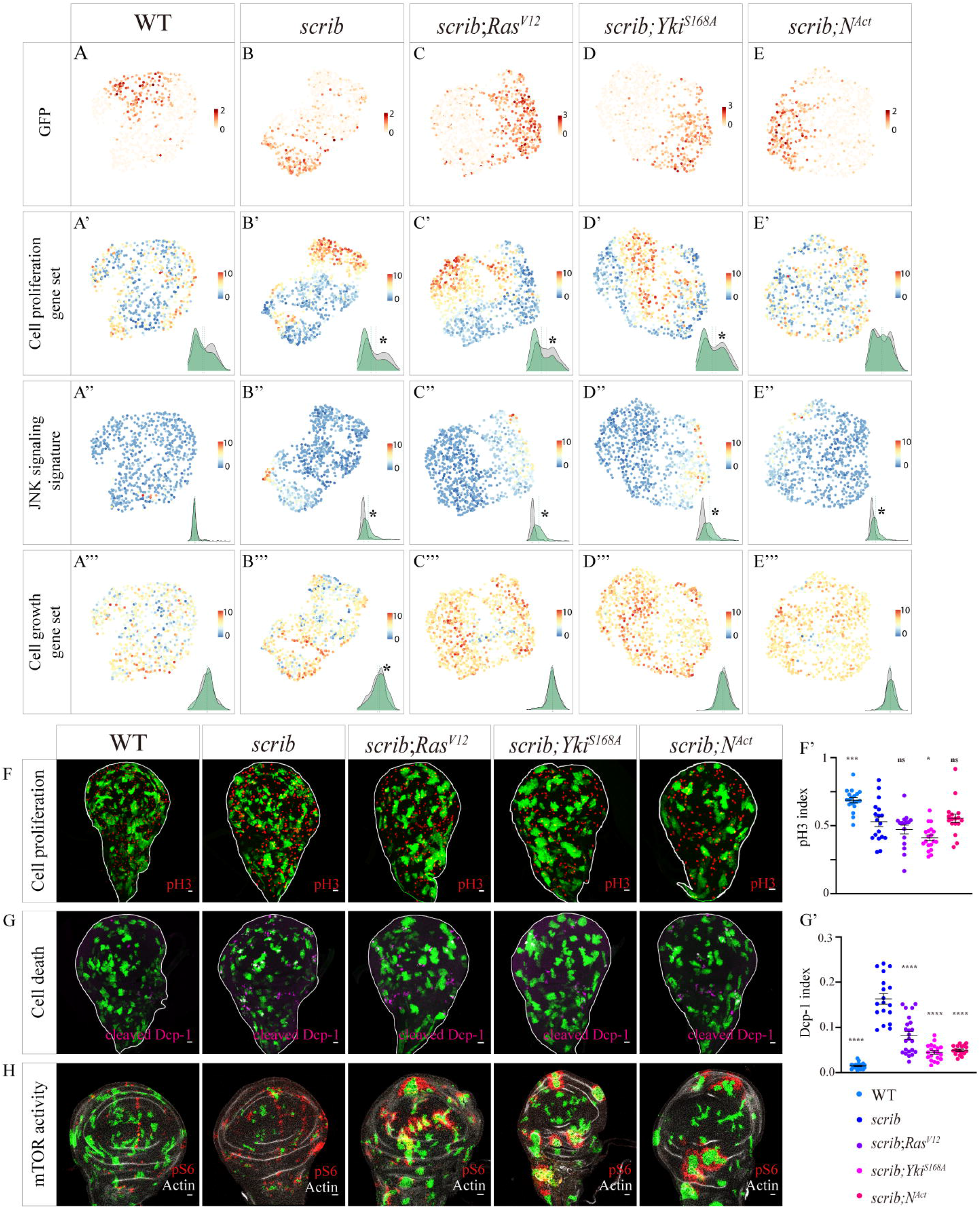
Analysis of proliferation, death and growth states of the *scrib*^-/-^ mutant, *scrib*^-/-^ *Ras^V12^*, *scrib*^-/-^ *Yki^S168A^* and *scrib*^-/-^ *N^Act^* clones. (A-E) UMAP projection of single cells from wing discs harboring MARCM clones of indicated genotypes. The cells are clustered using genes correlated with GFP gene expression for visualization. (A-E) are colored by the expression level of GFP or the expression score of respective gene sets. The insets are density plots of gene set expression score for GFP+ cells (colored in green) and GFP-cells (colored in gray). * p<0.001. Mann-Whitney test. The dash line represents the mean value of the gene set expression score. (F-H) Wing imaginal discs harboring MARCM clones of indicated genotypes inspected at 36 hours ACI are stained with anti-pH3 (red in F), anti-cleaved Dcp-1 (magenta in G) or anti-pS6 (red in H) antibodies. The MARCM clones are marked by the presence of GFP (green). Scale bar: 10μm. Quantification of pH3 Index and Dcp-1 Index in MARCM clones are in (F’) and (G’). ns p>0.05, *p<0.05, *** p<0.001, **** p<0.0001. One-way ANOVA test. Error bars represent mean with SEM.

The apoptosis of the *scrib* mutant clones is dependent on JNK signaling activation (*10*, *12*, *21*). We have previously identified a JNK signaling signature gene set composed of 106 genes (*22*)(Supplemental Table 3). We mapped the JNK signaling signature gene set to the current data and found that the GFP+ clonal cells exhibit elevated level of JNK target genes across all experimental groups except the control group (Figure 2A”-2E”). Mmp-1, a widely-used JNK signaling reporter, is indeed upregulated in the *scrib*^-/-^, *scrib*^-/-^ *Ras^V12^*, *scrib*^-/-^ *Yki^S168A^* and *scrib*^-/-^ *N^Act^* clones (Supplemental Figure 3). Interestingly, using Dcp-1 staining as a cell death marker, we found that the Dcp-1+ cell number is significantly reduced when Ras^V12^, Yki^S168A^ or N^Act^ is co-expressed in the *scrib* mutant clones (Figure 2G and 2G’). Together, these data suggest that Ras^V12^, Yki^S168A^ and N^Act^ signals can alleviate the *scrib* mutant clonal death without attenuating JNK signaling activation, consistent with previous findings that JNK signaling remains activated in the *scrib*^-/-^ *Ras^V12^* clones (*21*).

For cell growth state analysis, we have previously identified a cell growth gene set composed of 91 genes (*14*). Most of these genes are translation initiation factors, translation elongation factors and ribosomal genes (Supplemental Table 3). While the mechanism underlying high coexpression correlation of the 91 translation-related genes is unclear, we found that the expression of these genes is highly sensitive to insulin/mTOR signaling activity in transcriptomics datasets. Moreover, the expression of *thor*, a known mTOR signaling target, is very low in late larval imaginal discs (*23*) and only detected in a few cells in our single cell datasets. Therefore, we use this cell translation gene set as a proxy to analyze insulin/mTOR signaling activity from singlecell transcriptomics datasets. We note that in the *scrib* mutant clone group, there is a noticeable reduction of the cell growth gene set expression in the GFP-cells in comparison with the GFP+ *scrib* mutant clones (Figure 2B’”). The expression of the cell growth gene set is comparable in GFP+ and GFP-cells in all other groups (Figure 2A”’, C”’, D’”, E’“). We then use phospho-RpS6 (pS6) staining to monitor the mTOR signaling activity (*24*, *25*). Consistent with previous reports, in wing discs harboring control and the *scrib* mutant clones, the pS6 staining pattern is patchy (Figure 2H) (*24, 25*). Interestingly, in imaginal discs harboring the *scrib*^-/-^*Ras^V12^*, *scrib*^-/-^ *NICD* or *scrib*^-/-^*Yki^S168A^* clones, we observe a significant upregulation of pS6 signals in cells surrounding the GFP+ clones (Figure 2H). Together, these data suggest that Ras^V12^, Yki^S168A^ and N^Act^ signals can elevate mTOR signaling activity non-autonomously in cells adjacent to the *scrib* mutant clones.

### Upregulation of *upd2* in the *scrib*^-/-^ *Ras^V12^*, *scrib*^-/-^ *Yki^S168A^* and *scrib*^-/-^ *N^Act^* clones is necessary to promote tumor growth

Because Ras^V12^, Yki^S168A^ and N^Act^ signals can elevate mTOR signaling in the GFP-cells adjacent to the *scrib* mutant clones (Figure 2H), we compared GFP-wild type cells in all experimental groups (Figure 3A). Compared with GFP-cells in the *scrib* mutant group, the GFP-cells in the *scrib*^-/-^ *Ras^V12^*, *scrib*^-/-^ *Yki^S168A^* and *scrib*^-/-^ *N^Act^* groups have an overlap of 61 genes that are differentially expressed (Figure 3A) (Supplemental Table 4). Among the 61 common differentially-expressed genes, we noticed that the expression of *Socs36E* and *chinmo*, which are known targets of JAK/STAT signaling (*26*, *27*), are significantly upregulated in the GFP-cells of wing discs harboring the *scrib*^-/-^ *Ras^V12^*, *scrib*^-/-^ *Yki^S168A^* or *scrib*^-/-^ *N^Act^* clones in comparison with GFP-cells surrounding the *scrib* mutant clones (Figure 3B). Using 10xSTAT-GFP (*28*) as a reporter for JAK/STAT signaling activity, we found that the GFP-cells surrounding the *scrib^RNAi^ Ras^V12^*, *scrib^RNAi^ Yki^S168A^* or *scrib^RNAi^ N^Act^* clones frequently show elevated JAK/STAT signaling activity in comparison with GFP+ clones (Figure 3C). Moreover, when we generate clones that express a constitutively-active form of Stat92E (Stat92E^CA^)(*29*), we observe that mTOR signaling is frequently upregulated at the clonal boundary and within the Stat92E^CA^ clones (Figure 3D). Together these data suggest that JAK/STAT signaling activity is upregulated in GFP-wild type cells surrounding the *scrib*^-/-^ *Ras^V12^*, *scrib*^-/-^ *Yki^S168A^* or *scrib*^-/-^ *N^Act^* clones and JAK/STAT signaling discontinuity can lead to mTOR signaling activation at the signaling boundary.

**Figure 3.**
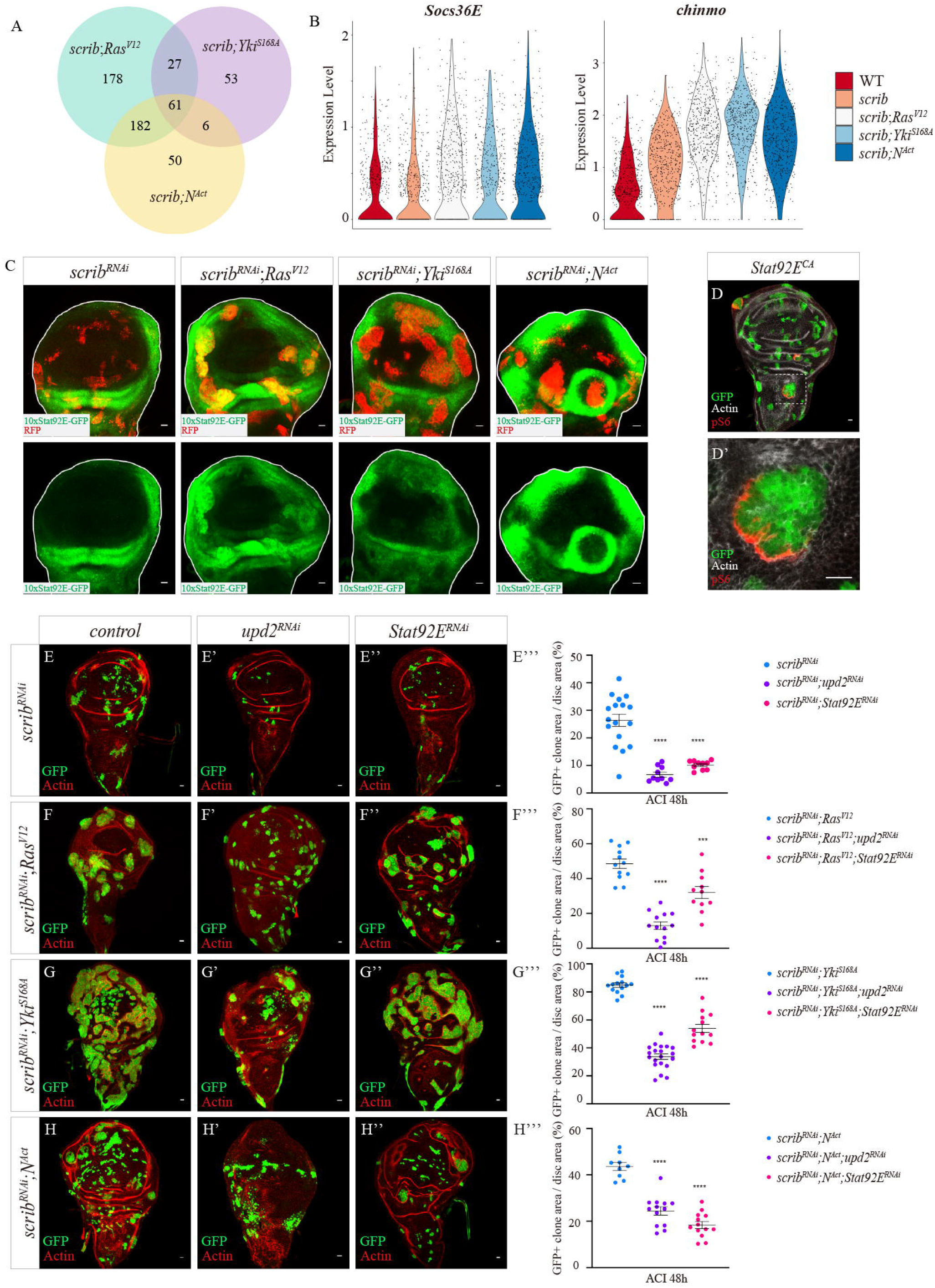
Upregulation of *upd2* in the *scrib*^-/-^ *Ras^V12^*, *scrib*^-/-^ *Yki^S168A^* and *scrib*^-/-^ *N^Act^* clones is necessary to promote tumor growth. (A) Venn diagram indicates the number of differentially expressed genes in the GFP-cells surrounding the *scrib*^-/-^ *Ras^V12^*, *scrib*^-/-^ *Yki^S168A^* and *scrib*^-/-^ *N^Act^* clones in comparison with the GFP-cells surrounding the *scrib* mutant clones. (B) Violin plot of *Socs36E* and *chinmo* expression level in the GFP-cells from wing discs harboring MARCM clones of indicated genotypes. (C) JAK/STAT signaling activity is indicated by *10xStat92E-GFP* reporter (green) from wing discs harboring clones of indicated genotypes at 48 hours ACI. The clones are marked by the presence of RFP (red). Scale bar: 10μm. (D-D’) Wing imaginal discs harboring the *Stat92ECA* clones inspected at 48 hours ACI are stained with anti-pS6 antibody (red). The clones are marked by the presence of GFP (green). (D’) is a zoom-in image of the region highlighted by dotted line in (D). Note that pS6 staining is within the *Stat92ECA* clones. Scale bar: 10μm. (E-H) Wing imaginal discs harboring FlpOut clones of indicated genotypes are inspected at 48hours ACI. The clones are marked by the presence of GFP (green) and imaginal discs are stained with phalloidin to visualize cell outline (red). Scale bar: 10μm. Quantification of the percentage of GFP+ clone area in wing discs is shown in (E’”-H”‘). *** p<0.001, **** p<0.0001. One-way ANOVA test. Error bars represent mean with SEM.

To identify the source of JAK/STAT signaling activation in GFP-cells surrounding the *scrib*^-/-^ *Ras^V12^*, *scrib*^-/-^ *Yki^S168A^* and *scrib*^-/-^ *N^Act^* clones, we examined the expression of Unpaired family cytokines, which are known ligands that activate JAK/STAT signaling (*30*–*33*). We found that *upd2* is upregulated in *scrib^RNAi^ Ras^V12^*, *scrib^RNAi^ Yki^S168A^* and *scrib^RNAi^ N^Act^* clones at this stage through mRNA FISH experiments (Supplemental Figure 4). Interestingly, when we depleted Upd2 in the *scrib^RNAi^ Ras^V12^*, *scrib^RNAi^ Yki^S168A^* or *scrib^RNAi^ N^Act^* clones using RNAi, the size of these clones is significantly reduced (Figure 3E-3H, 3E’-3H’). We adopted a sgRNA construct that knocks out *upd1-3* simultaneously and found that removal of Upd1-3 also has a strong effect on repressing the *scrib*^-/-^ *Yki^S168A^* and *scrib*^-/-^ *N^Act^* clonal tumor growth (Supplemental figure 5).

JAK/STAT signaling activity is known to affect the *scrib* and *dlg* mutant growth (*34*, *35*). It was shown that expression of a dominant negative form of Domeless receptor (Dome^DN^) can suppress the growth of *scrib*^-/-^ *Ras^V12^* tumors and co-expression of *Ras^V12^* and *Upd2* lead to invasive tumor growth (*34*). The expression of *Dome^DN^* can also attenuate the growth of *scrib^RNAi^* tumors driven with a pan-wing disc driver (*35*). To examine whether the effects of *upd2* RNAi is due to JAK/STAT signaling attenuation within these clones, we depleted Stat92E using RNAi (*36*–*38*) and found that the clonal growth is indeed reduced (Figure 3E”-3H”). However, for the Ras and Yki groups, the reduction of tumorous growth using *Stat92E* RNAi is overall weaker than the effects of *upd2* RNAi (Figure 3F’”-3G’”). Moreover, the co-expression of Stat92E^CA^ in the *scrib* mutant clones does not promote tumorous growth to the same degree as Yki, Ras and Notch signals, except at the hinge region, consistent with previous reports (Supplemental Figure 6)(*39*). Together, our data are consistent with previous findings that JAK/STAT signaling activity is important for determining the growth outcome of the *scrib* mutant clones. However, our data show that JAK/STAT signaling activity is strongly activated in wild type cells surrounding the *scrib^RNAi^ Ras^V12^*, *scrib^RNAi^ Yki^S168A^* and *scrib^RNAi^ N^Act^* clones, which suggest that JAK/STAT activation in wild type neighboring cells may also contribute to the clonal tumor growth.

### Upd2 is a cell competition factor in wing discs

As a control experiment, we generated *upd2* RNAi clones or *upd1-3* sgRNA-Cas9 clones in wing discs and found that *upd2* RNAi clones or *upd1-3* sgRNA-Cas9 clones cannot survive in wing discs (Figure 4A-B) (Supplemental Figure 7). The result is unexpected because *upd2* loss-of-function mutants are viable and give rise to normal wing discs (*30*, *32*). Moreover, when we use *nubbin-Gal4* to drive the same *upd2* RNAi construct, the size of wing pouch has no significant difference in comparison with control (Figure 4E–4F). Therefore, the elimination of *upd2* RNAi clones closely resembles the cell competition process in that viable clonal cells are eliminated when they encounter neighboring cells of different genotypes.

**Figure 4.**
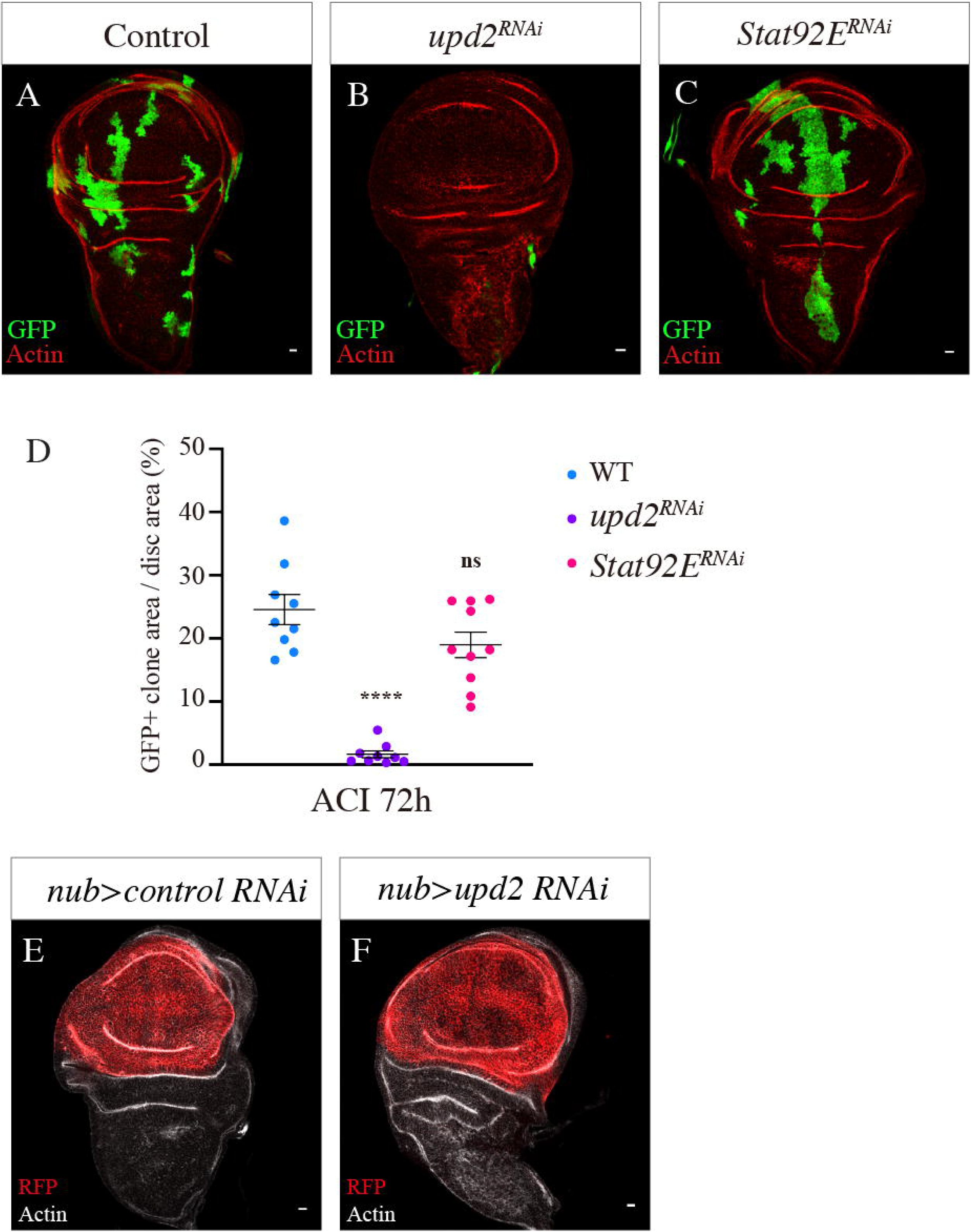
Upd2 is a cell competition factor in wing discs. (A-D) Wing imaginal discs harboring FlpOut clones of indicated genotypes are inspected at 72hours ACI. The clones are marked by the presence of GFP (green) and imaginal discs are stained with phalloidin to visualize cell outline (red). Scale bar: 10μm. Quantification of the percentage of GFP+ clone area in wing discs is shown in D. ns p>0.05, **** p<0.0001. One-way ANOVA test. Error bars represent mean with SEM. (E-F) Confocal images of wing imaginal discs of *upd2* knockdown using a uniform wing pouch driver. Imaginal discs are stained with phalloidin to visualize cell outline (white). Scale bar: 10μm.

JAK/STAT signaling has long been known to play an important role in the cell competition process (*40*–*44*). For example, in wing discs, the *Stat92E* mutant clones do not survive well when the clones are induced early during disc development (*44*). When *Dome^DN^* is expressed in the midgut, the expansion of wild type clones in *Minute* background are reduced (*43*). Similarly, it was shown that JAK/STAT signaling activity is required for competition-induced winner cell expansion (*42*). In the fly testis, the *socs36E* mutant cyst progenitor cell (CPC) clones outcompete other CPCs (*41*). Similarly, the *chinmo* mutant germline stem cell (GSC) clones outcompete other GSCs (*40*). Consistent with these findings, our data support a general involvement of JAK/STAT signaling in cell competition. However, we found that *Stat92E* RNAi clones survive well in wing discs in comparison with a strong elimination of *upd2* RNAi clones (Figure 4B–4D), indicating that the elimination of *upd2* RNAi or *upd1-3* sgRNA-Cas9 clones may not be due to cell-autonomous reduction of JAK/STAT activity.

Ras, YAP and Notch oncogenic signals are frequent drivers of tumorigenesis in human cancers. Like human cancers, the fly tumors also exhibit a strong degree of robustness in that different oncogenic signals are interchangeable in driving tumor formation. Our study here demonstrated that oncogenic Ras^V12^, Yki^S168A^ and N^Act^ signals converge to upregulate Upd2 and promote the *scrib* mutant clonal survival. While *Upd2* homozygous mutant wing discs are completely normal, the Upd2-deficient clones undergo elimination when they are surrounded by wildtype cells. Therefore, we identified Upd2 as a cell competition factor in *Drosophila* wing imaginal discs.

## Materials and methods

### Drosophila stocks and Genetics

Drosophila melanogaster strains were listed in Supplemental Table 5. All experimental crosses were raised at 25°C in uncrowded conditions. To generate MARCM clones, around 50 embryos were collected within 3 hours and raised until 60 hours after egg laying (AEL). The larvae were then placed in water bath for 1 hour at 37°C for heat shock and dissected at 36 hours or 60 hours after clone induction (ACI) for clone inspection. To generate FlpOut clones, the larvae were raised until 72 hours AEL and then placed in water bath for 15min at 37°C for heat shock and dissected at 48 hours or 72 hours ACI for clone inspection.

### Generation of transgenic flies

Full length of Yki DNA sequence was synthesized, and the point mutation on S168A site was introduced using primers listed in Supplemental Table 5. The resulting Yki^S168A^ fragment was cloned into pTiger (*45*), sequence verified and targeted to attP site at 25C6 and 2A3, respectively (WellGenetics). The NICD construct in this study was cloned from UAS-NICD strain (*46*), in which the Notch extracellular domain including the signal peptide and transmembrane domain was deleted. The cloned NICD fragment was then cloned into pTiger, sequence verified and targeted to attP sites at 68A4 and at 19E7, respectively (WellGenetics).

### Immunofluorescence and microscopy

Larvae were dissected in 1xPBS, fixed with 4% PFA for 20 minutes, and blocked with 1% BSA in 0.1%PBST for 1 hour at room temperature. The larvae were then incubated with primary antibodies overnight at 4°C and secondary antibodies for 2 hours at room temperature. The primary and secondary antibodies were listed in Supplemental Table 5. Phalloidin conjugated with Alexa Fluor dyes (1:1000, Thermo Fisher Scientific) and Hoechst 33342 (1:1000, Thermo Fisher Scientific) were used to stain F-actin and DNA, respectively. Images were taken with a Leica TCS SP8 confocal microscope as z-stacks with a step size of 1μm.

### Fluorescent RNA in situ hybridization (FISH)

*Upd2* FISH were performed following Hybridzation chain reaction (HCR) v3.0 protocol(*47*). Probe sets against upd2 mRNA ordered from BGI Tech company were listed in Supplemental Table 5.

### Image Processing Analysis and Quantification

Each image stack is converted into a merged image by using “Max Intensity” function of Fiji (version “2.3.0”). The merged image was split into multiple channels. We loaded the DNA channel image into CellProfiler (version “4.0.1”) environment and applied “Gaussian filter” to smooth the image. The smoothed image was piped into “IdentifyPrimaryObjects” module to identify wing imaginal discs by setting the diameter of objects from 500 to 2000 pixels and applying Sauvola thresholding method. Using the GFP channel image, we identified GFP clones by setting the diameter of objects from 5 to 500 pixels and applying Otsu thresholding method in “IdentifyPrimaryObjects” module. To identify pH3 or Dcp-1 signals, we set the diameter of objects from 4 to 40 pixels and used Otsu thresholding method in “IdentifyPrimaryObjects” module. When assigning the signals of PH3 and Dcp-1 into the clones, we used “RelateObjects” module to determine the signals inside the clones. Using pixel as unit, we measured the area of these objects by adopting “MeasureImageAraeOccupied” module.

Statistical analysis was performed with Excel and Prism 9. The unpaired T-test was performed for two independent variables that are normally distributed with no significant differences in variance. The one-way ANOVA test was performed for three or more independent variables that are normally distributed with no significant differences in variance. For all plots, the error bar indicates the mean with SEM. P values are presented as: ≥ 0.05 (ns), 0.01-0.05 (*), 0.001-0.01 (**), 0.0001-0.001 (***), < 0.0001 (****) unless otherwise specified.

### 10x Genomics Single-cell RNA-sequencing and data analysis

Wing imaginal discs were dissected in Dulbecco’s phosphate-buffered saline (DPBS; ThermoFisher) and dissociated in 0.25% trypsin-EDTA solution at 37°C for 10 min. Cells were then washed by DPBS twice and passed through a 35μm filter before library preparation. Construction of 10x single-cell libraries and sequencing on the Illumina Hiseq platform were performed by Novogene.

Raw data mapping was performed by standard Cell Ranger pipeline (v 2.2.0) to generate UMI count matrices. Reads alignment was based on the BDGP6 genome reference fastaq file and annotated by the BDGP6.91.gtf file. The UMI count matrices were loaded into R environment to build Seurat (v4.3.0) objects. To reduce the noise of the dataset, we filtered out cells with UMI counts more than 50000 and cells that express less than 1500 genes; we also removed genes detected in less than 10 cells. The filtered dataset was then used for downstream analysis. Expression data was normalized using the default LogNormalize and SCTransform functions in Seurat. Variable genes were selected using the Seurat FindVariableGenes function. Dimension reduction, cell clustering, and cell marker identification were conducted with default parameters. Since the numbers of GFP+ cells were low, all GFP+ cells and 500 randomly selected GFP-cells from each genotype were used in subsequent analysis. To conduct cross-sample analysis, we removed batch-effect genes we identified from two *scrib* mutant samples sequenced in two batch of experiments. To compare the expression level of single gene across samples, we used VlnPlot function for visualization and applied non-parametric Kruskal-Wallis test to examine the statistical significance. For visualize GFP+ and GFP-cells in UMAP projection, we calculated the Spearman correlation coefficients between normalized gene expression values of GFP and other genes by using the cor function in R. Genes with correlation coefficient greater than 0.15 were retained as variable genes to project GFP+ cells from GFP-cells. To quantify the activity of a specific gene set, for each gene inside the gene set, we first calculated z-score based on log-normalized expression values of all cells and then computed the mean value of the z-scores from all the genes in this gene set, which was used as the expression score of this gene set. To visualize the expression score, we added the expression score data into the metadata file of Seurat project and applied FeaturePlot function to project the expression score data to UMAP plot. The gene sets used to generate projections are manually curated from FlyBase and existing literatures. To compare gene set activities between GFP+ and GFP-cells, the expression score of a specific gene set was plotted as a density plot using geom_density function. Nonparametric Mann-Whitney test was performed to test the statistical significance.

## Acknowledgments

We would like to thank Drs. Mark Van Doren, Jose Pastor-Pareja, Erika Bach, Aurelio Teleman for kindly sharing fly strains and antibodies. This work was supported by grants to Y. Yan from HKRGC/GRF16103620 and Shenzhen Science and Technology Innovation Commission/JCYJ20200109140201722.

## Author Contributions

Y.Wang and Y. Yan designed the research and wrote the manuscript; Y.Wang conducted the experiments; J. J. He and M.X. Deng performed the data analysis.

