## Supplementary figures and images for "Oncogenic Ras, Yki and Notch signals converge to confer clone competitiveness through Upd2"

### Supplemental Figure 1

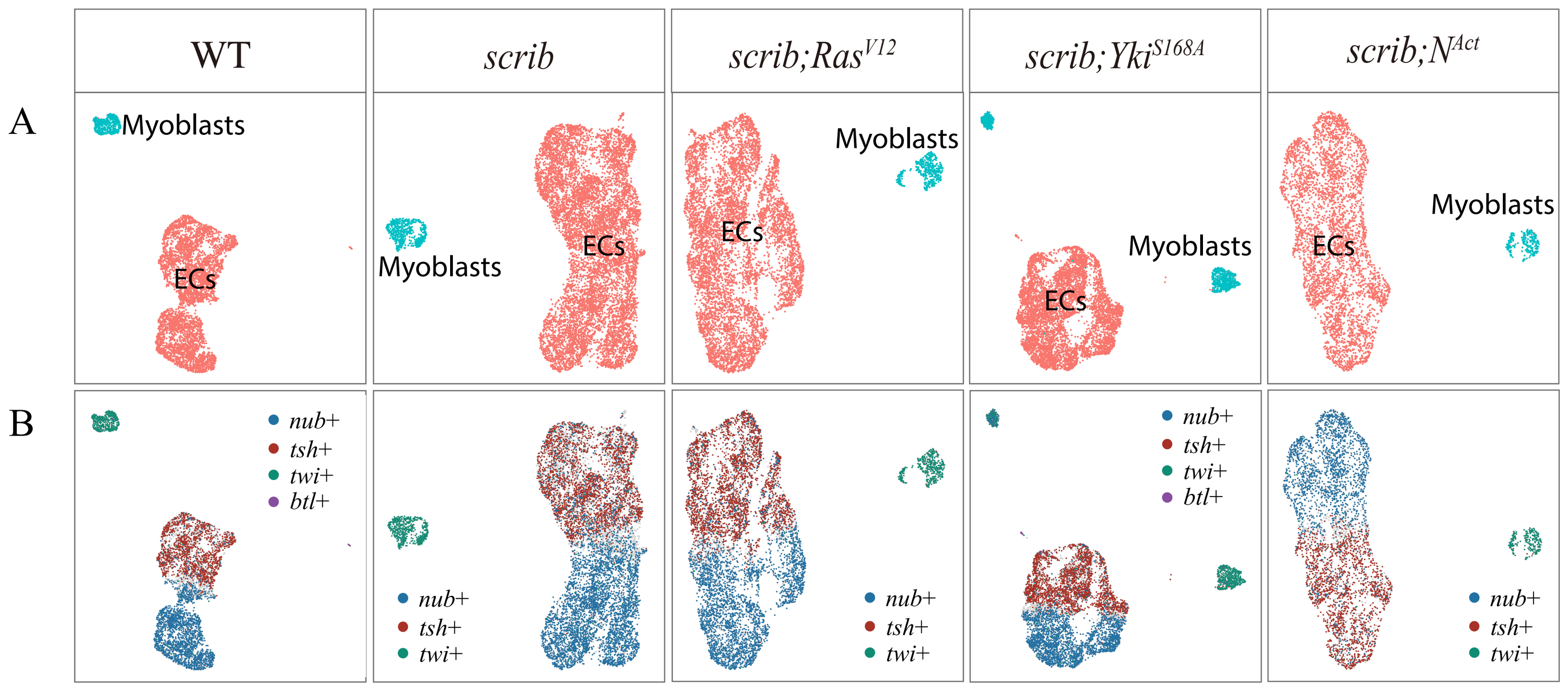

### Supplemental Figure 2

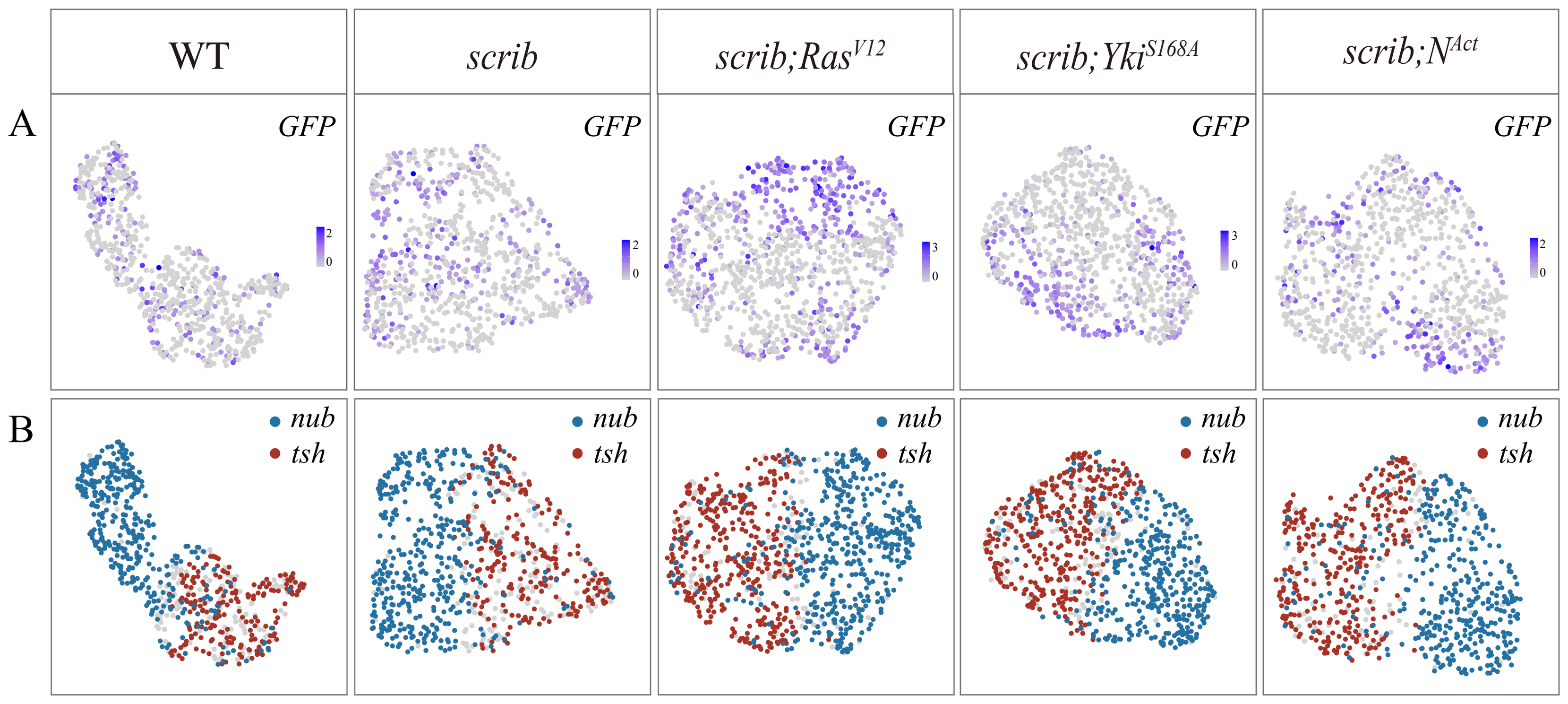

### Supplemental Figure 3

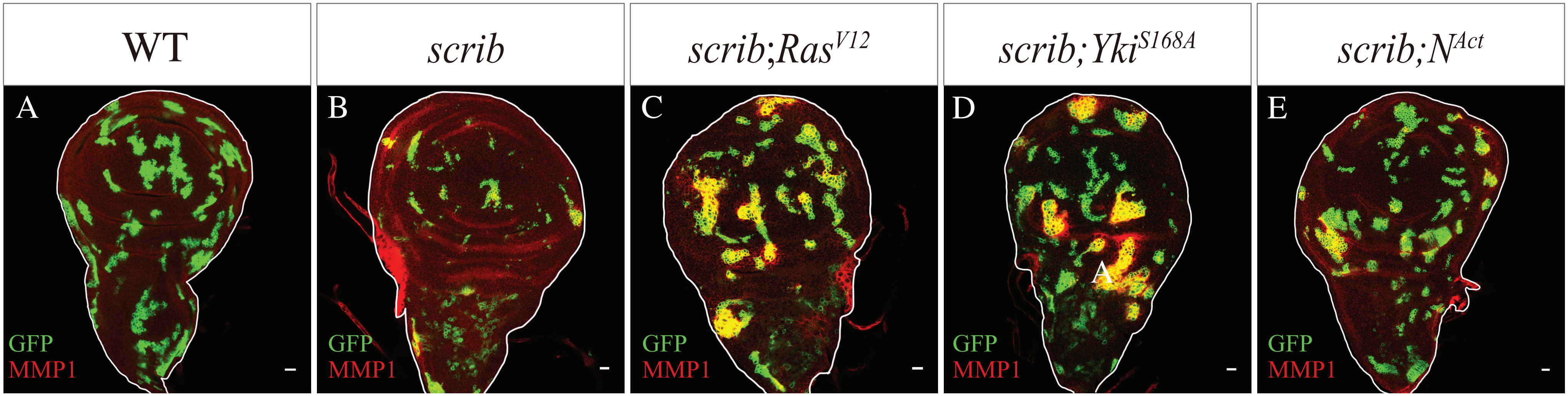

### Supplemental Figure 4

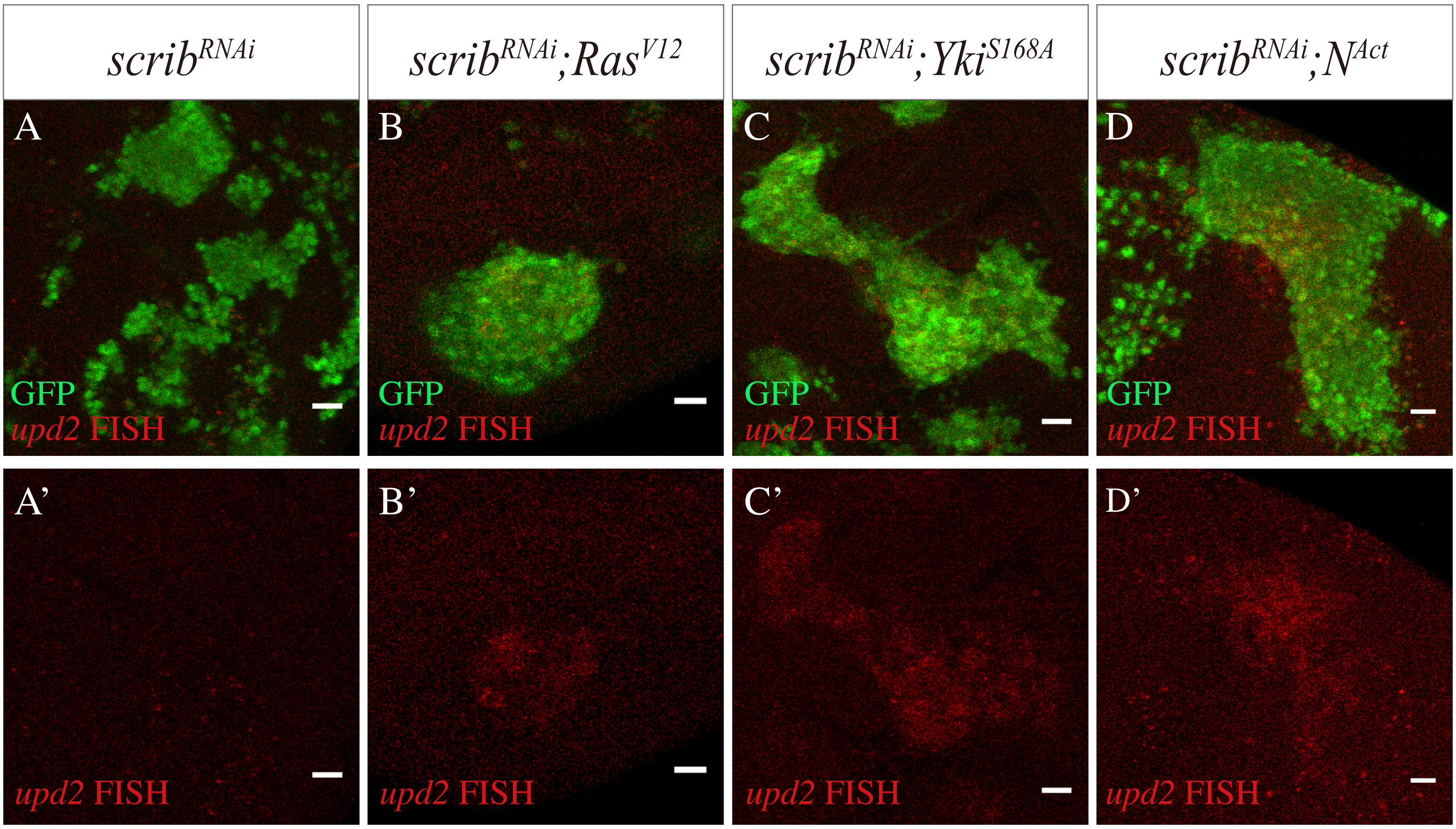

### Supplemental Figure 5

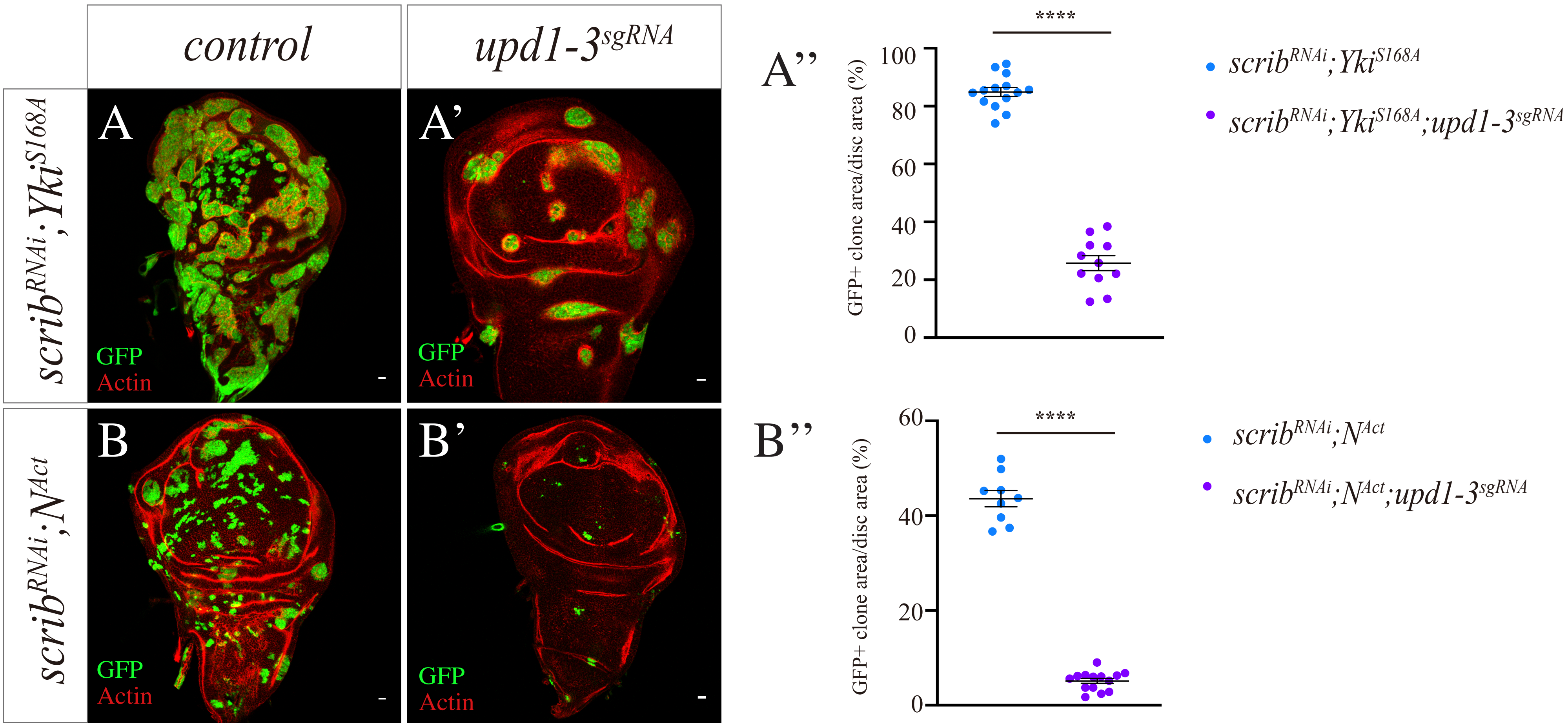

### Supplemental Figure 6

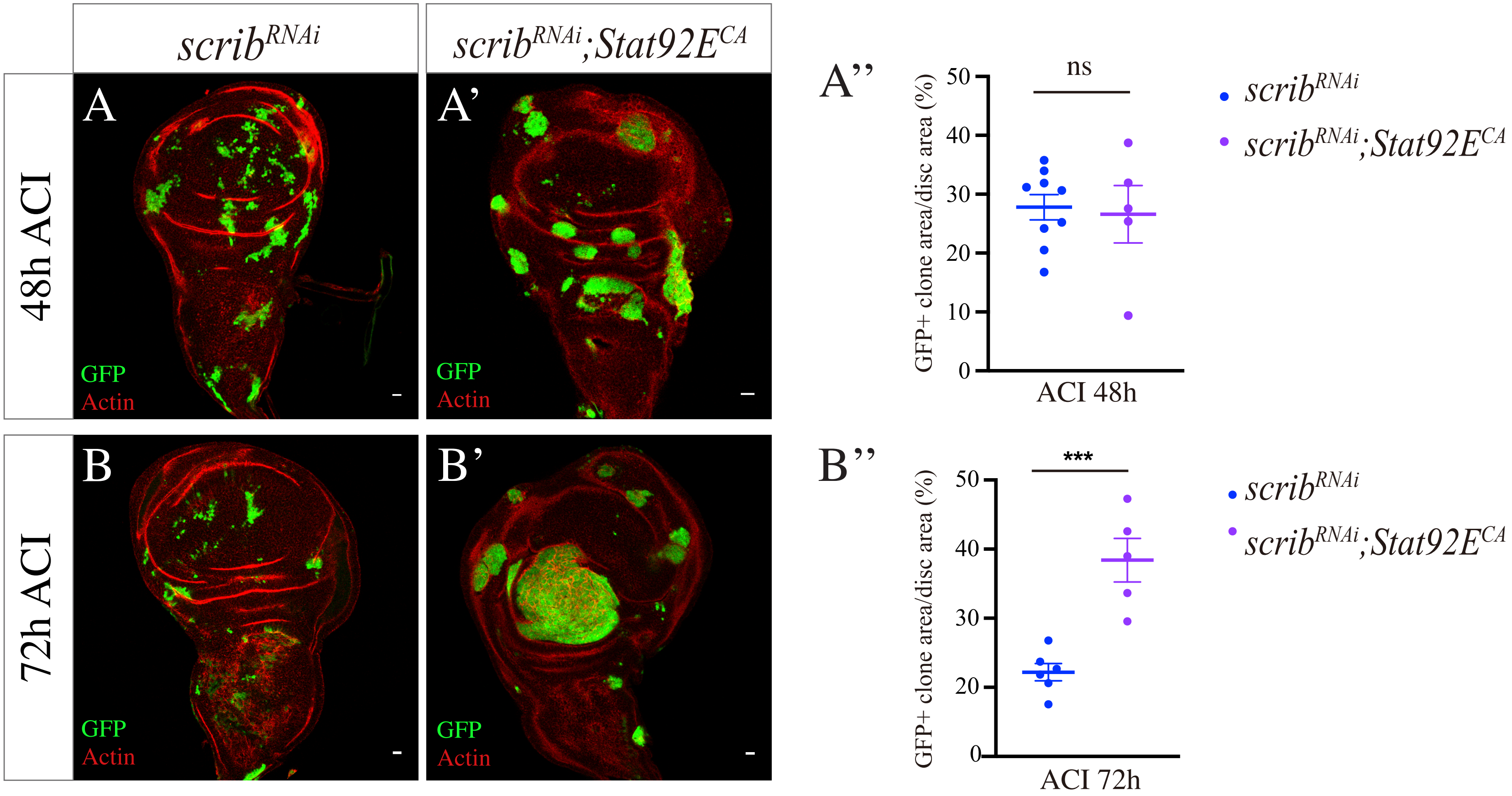

### Supplemental Figure 7

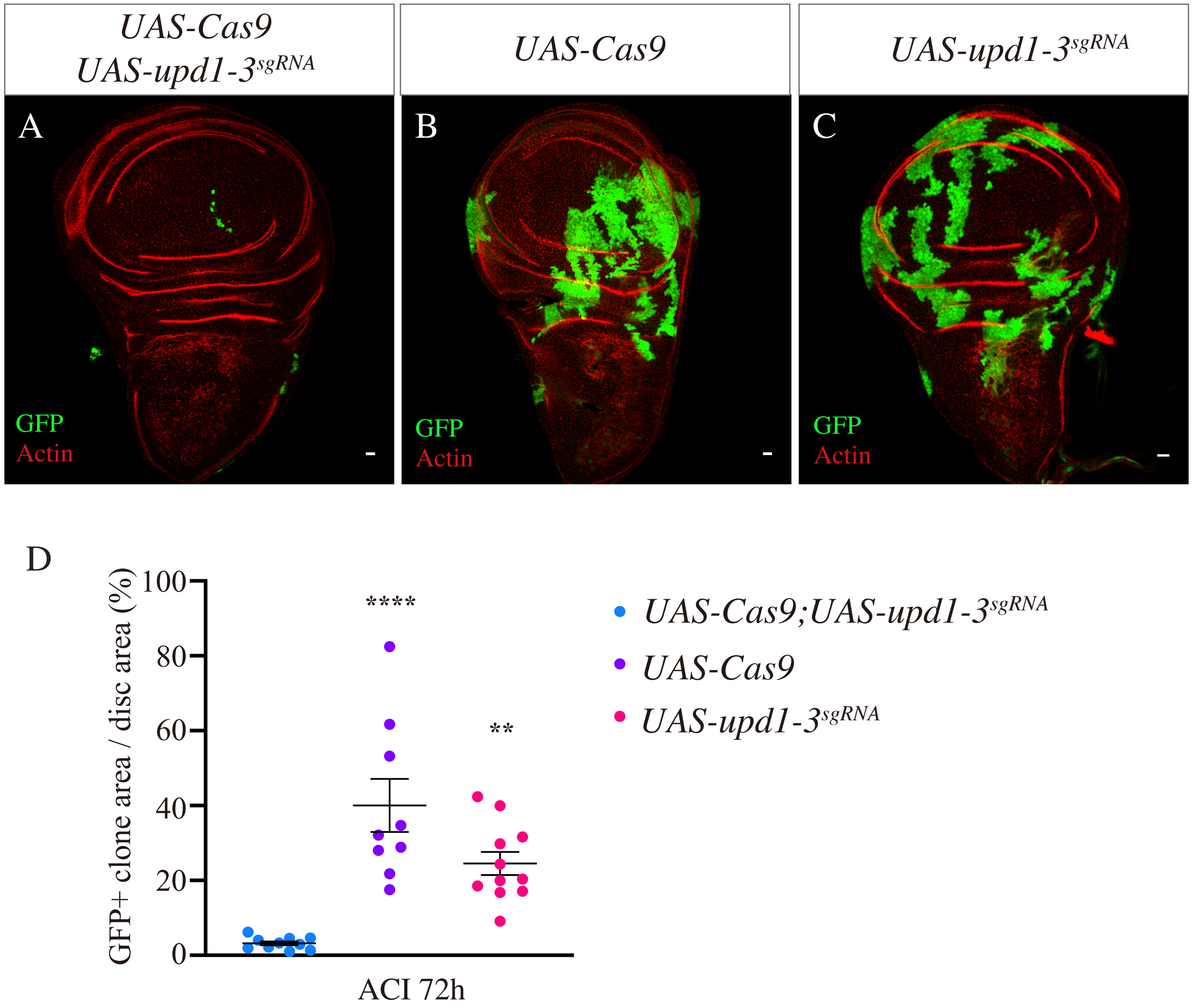
