## Supplemental Figure legend for "Oncogenic Ras, Yki and Notch signals converge to confer clone competitiveness through Upd2"

**Supplemental Figure 1 Overview of single cells from wing discs harboring MARCM clones of indicated genotypes.**

(A) UMAP projection of single cells labeled by cell types. ECs: wing disc epithelial cells.

(B) UMAP projection of single cells labelled by marker genes. *Twist* (*twi*) is specifically expressed in myoblasts. *Breathless* (*btl*) labels tracheal cells. *Nubbin* (*nub*) and *teashirt* (*tsh*) are proximal-distal identity markers for wing disc epithelial cells.

**Supplemental Figure 2 Random selection of 500 GFP- cells to balance the GFP+ and GFP- cell number does not affect the major clustering feature.**

(A) UMAP projection of all GFP+ and 500 randomly selected GFP- epithelial cells colored by the expression level of *GFP* from wing discs harboring MARCM clones of indicated genotypes.

(B) UMAP projection of single cells labelled by *nubbin* (*nub*) and *teashirt* (*tsh*).

**Supplemental Figure 3 The *scrib*^-/-^, *scrib^-/-^* *Ras^V12^, scrib^-/-^* *Yki^S168A^* and *scrib^-/-^* *N^Act^* clones show elevated JNK signaling activity.**

(A-E) Wing imaginal discs harboring MARCM clones of indicated genotypes (48 hours ACI) are stained with anti-MMP1 antibody (red). The MARCM clones are marked by the presence of GFP (green). Scale bar: 10μm.

**Supplemental Figure 4 The *scrib^-/-^* *Ras^V12^, scrib^-/-^* *Yki^S168A^* and *scrib^-/-^* *N^Act^* clones elevate *upd2* transcription.**

(A-E) Confocal images of *upd2* mRNA FISH (red) from wing imaginal discs harboring FlpOut clones of indicated genotypes. The clones are marked by the presence of GFP (green). Scale bar: 10μm.

**Supplemental Figure 5 Removal of Upd 1-3 blocks *scrib^RNAi^ Yki^S168A^* and *scrib^-/-^* *N^Act^* clonal growth.**

(A- B) Confocal images of wing imaginal discs harboring FlpOut clones of indicated genotypes inspected at 48 hours ACI. The clones are marked by the presence of GFP (green) and imaginal discs are stained with phalloidin to visualize cell outline (red). Scale bar: 10μm. Quantification of the percentage of GFP+ clone area in wing discs is shown in (A’’) and (B’’). **** p<0.0001. Unpaired t test. Error bars represent mean with SEM.

**Supplemental Figure 6 Stat^CA^ can promote the *scrib^RNAi^* clone growth at the hinge region.**

(A-B) Confocal images of wing imaginal discs harboring FlpOut clones of indicated genotypes inspected at 48 hours ACI (A-A’) and 72 hours ACI (B-B’). The clones are marked by the presence of GFP (green) and imaginal discs are stained with phalloidin to visualize cell outline (red). Scale bar: 10μm. Quantification of the percentage of GFP+ clone area in wing discs is shown in (A’’) and (B’’). ns p>0.05, *** p<0.001. Unpaired t test. Error bars represent mean with SEM.

**Supplemental Figure 7 Removal of Upd 1-3 lead to clonal elimination.**

(A-C) Confocal images of wing imaginal discs harboring FlpOut clones of indicated genotypes inspected at 72 hours ACI. The clones are marked by the presence of GFP (green) and imaginal discs are stained with phalloidin to visualize cell outline (red). Scale bar: 10μm. Quantification of the percentage of GFP+ clone area in wing discs is shown in (D). ** p<0.01, **** p<0.0001. One-way ANOVA test. Error bars represent mean with SEM.

**Supplemental Table 1: Summary of 10x single-cell RNA sequencing results.**

**Supplemental Table 2: Gene expression correlation coefficient with GFP**

**Supplemental Table 3: Curated gene set list.**

**Supplemental Table 4: List of differentially expressed genes in GFP- cells.**

**Supplemental Table 5: Key Resources Table.**

**Drosophila genotypes by figure**

**Figure 1-2, Supplemental Figure 1-3**

**WT**

*hsFlp; tub>y+>Gal4 UAS-GFP/+; tub-Gal80 FRT82B/FRT82B*

***scrib***

*hsFlp; tub>y+>Gal4 UAS-GFP/+; tub-Gal80 FRT82B/scrib^1^FRT82B*

***scrib;Ras^V12^***

*hsFlp; tub>y+>Gal4 UAS-GFP/UAS-Ras^V12^; tub-Gal80 FRT82B/scrib^1^FRT82B*

***scrib;Yki^S168A^***

*hsFlp; tub>y+>Gal4 UAS-GFP/UAS-Yki^S168A^; tub-Gal80 FRT82B/scrib^1^FRT82B*

***scrib;N^Act^***

*hsFlp; tub>y+>Gal4 UAS-GFP/UAS-N^Act^; tub-Gal80 FRT82B/scrib^1^FRT82B*

**Figure 3:**

(C)

***scrib^RNAi^***

*hsFlp; act>y+>Gal4 UAS-mRFP, UAS-scrib RNAi/10XStat92E-GFP*

***scrib^RNAi^;Ras^V12^***

*hsFlp;act>y+>Gal4 UAS-mRFP, UAS-scrib RNAi/10XStat92E-GFP;*

*UAS-Ras^V12^/+*

***scrib^RNAi^;Yki^S168A^***

*hsFlp;act>y+>Gal4 UAS-mRFP, UAS-scrib RNAi/10XStat92E-GFP;*

*UAS-Yki^S168A^/+*

***scrib^RNAi^;N^Act^***

*hsFlp;act>y+>Gal4 UAS-RFP, UAS-scrib RNAi/10XStat92E-GFP;*

*UAS-N^Act^/+*

(D) *hsFlp; act>y+>Gal4 UAS-GFP/ UAS-3HA-Stat92E^ΔNΔC^；UAS-3HA- Stat92E^ΔNΔC^*／+

(E) *hsFlp; act>y+>Gal4 UAS-GFP, UAS-scrib RNAi/+*

(E’) *hsFlp; act>y+>Gal4 UAS-GFP, UAS-scrib RNAi/+; UAS-upd2 RNAi/+*

(E’’) *hsFlp; act>y+>Gal4 UAS-GFP, UAS-scrib RNAi/+; UAS-Stat92E RNAi/+*

(F) *hsFlp; act>y+>Gal4 UAS-GFP, UAS-scrib RNAi/UAS-Ras^V12^*

(F’) *hsFlp; act>y+>Gal4 UAS-GFP, UAS-scrib RNAi/UAS-Ras^V12^; UAS-upd2 RNAi/+*

(F’’) *hsFlp; act>y+>Gal4 UAS-GFP, UAS-scrib RNAi/UAS-Ras^V12^; UAS- Stat92E RNAi/+*

(G) *hsFlp, UAS-Yki^S168A^; act>y+>Gal4 UAS-GFP, UAS-scrib RNAi/+*

(G’) *hsFlp, UAS-Yki^S168A^; act>y+>Gal4 UAS-GFP, UAS-scrib RNAi/+; UAS-upd2 RNAi/+*

(G’’) *hsFlp, UAS-Yki^S168A^; act>y+>Gal4 UAS-GFP, UAS-scrib RNAi/+; UAS- Stat92E RNAi/+*

(H) *hsFlp, UAS-N^Act^; act>y+>Gal4 UAS-GFP, UAS-scrib RNAi/+*

(H’) *hsFlp, UAS- N^Act^; act>y+>Gal4 UAS-GFP, UAS-scrib RNAi/+; UAS-upd2 RNAi/+*

(H’’) *hsFlp, UAS- N^Act^; Act>y+>Gal4 UAS-GFP, UAS-scrib RNAi/+; UAS- Stat92E RNAi/+*

**Figure 4:**

(A) *hsFlp; act>y+>Gal4 UAS-GFP/+; UAS-36303 RNAi/+*

(B) *hsFlp; act>y+>Gal4 UAS-GFP/+; UAS-upd2 RNAi/+*

(C) *hsFlp; act>y+>Gal4 UAS-GFP/+; UAS-Stat92E RNAi/+*

(F) *nubbin-Gal4 UAS-RFP/+; UAS-36303 RNAi/+*

(G) *nubbin-Gal4 UAS-RFP/+; UAS-upd2 RNAi/+*

**Supplemental Figure 4**

(A) *hsFlp; act>y+>Gal4 UAS-GFP, UAS-scrib RNAi/+*
(B) *hsFlp; act>y+>Gal4 UAS-GFP, UAS-scrib RNAi/UAS-Ras^V12^*
(C) *hsFlp; act>y+>Gal4 UAS-GFP, UAS-scrib RNAi/UAS-Yki^S168A^*
(D) *hsFlp; act>y+>Gal4 UAS-GFP, UAS-scrib RNAi/UAS-N^Act^*

**Supplemental Figure 5**

(A) *hsFlp, UAS-Yki^S168A^; act>y+>Gal4 UAS-GFP, UAS-scrib RNAi/+*

(A’) *hsFlp, UAS-Yki^S168A^; act>y+>Gal4 UAS-GFP, UAS-scrib RNAi/ upd1-3^sgRNA^; UAS-Cas9/+*

(B) *hsFlp, UAS-N^Act^; act>y+>Gal4 UAS-GFP, UAS-scrib RNAi/+*

(B’) *hsFlp, UAS-N^Act^; act>y+>Gal4 UAS-GFP, UAS-scrib RNAi/ upd1-3^sgRNA^; UAS-Cas9/+*

**Supplemental Figure 6**

(A) and (B) *hsFlp; act>y+>Gal4 UAS-GFP, UAS-scrib RNAi/+*

(A’) and (B’) *hsFlp; act>y+>Gal4 UAS-GFP, UAS-scrib RNAi/ UAS-3HA-Stat92E^ΔNΔC^; UAS-3HA-Stat92E^ΔNΔC^/+*

**Supplemental Figure 7**

(A) *hsFlp; act>y+>Gal4 UAS-GFP/ upd1-3^sgRNA^; UAS-Cas9/+*

(B) *hsFlp; act>y+>Gal4 UAS-GFP/+; UAS-Cas9/+*

(C) *hsFlp; act>y+>Gal4 UAS-GFP/ upd1-3^sgRNA^*
